# Insights into the Datasets, Tools, and Training Needs of the AnVIL Community: 2024

**DOI:** 10.1101/2025.11.14.688517

**Authors:** Kathryn J. Isaac, Katherine E. L. Cox, Kai Yin Ho, Elizabeth M. Humphries, Natalie Kucher, Jeffrey T. Leek, Stephen Mosher, Michael C. Schatz, Frederick J. Tan, Ava M. Hoffman

**Affiliations:** Fred Hutch Cancer Center, Biostatistics Program, Seattle, WA, USA; Johns Hopkins University, Department of Biology, Baltimore, MD, USA; Vanderbilt University Medical Center, Nashville, TN, USA; Johns Hopkins University, Department of Biology, Baltimore, MD, USA and Johns Hopkins University, Department of Computer Science, Baltimore, MD, USA; Fred Hutch Cancer Center, Biostatistics Program, Seattle, WA, USA and Johns Hopkins Bloomberg School of Public Health, Department of Biostatistics, Baltimore, MD, USA

## Abstract

The NHGRI Genomic Data Science Analysis, Visualization, and Informatics Lab-space (AnVIL) provides a secure cloud-based environment where research and education communities can analyze genomic and biomedical data. The platform supports a wide range of data analysis as well as the ability to safely store and access data in compliance with NIH policies. Work on the AnVIL platform can be easily shared to promote reproducible science and collaboration. The purpose of this study is to better understand the current user base of the AnVIL platform. The AnVIL Community Poll aimed to collect baseline information, identify development opportunities, guide the prioritization of user support strategies, and succinctly but comprehensively describe the current AnVIL Community. The AnVIL Team disseminated the inaugural AnVIL Community Poll by sharing it broadly on social media and through AnVIL and related consortia mailing lists. We categorized respondents as either returning or potential users of the AnVIL platform (based on their provided usage description) and examined user experiences: specifically user backgrounds, technological comfort, research interests, computational needs, and preferences for training and support. Our sample of the AnVIL community found opportunities for platform adoption beyond the current user base and identified areas where training should be enhanced, training preferences, and user computational needs. Specifically, while most respondents were involved in human genomics research, there may be potential for growth in adoption of the platform by prioritizing materials to support clinical researchers. All respondents felt availability of specific tools or datasets was a key feature of the platform. The broader community may also benefit from further development or showcasing of resources to facilitate cost management, finding and incorporating analysis tools, and data import. Our sample greatly preferred virtual training opportunities and returning users of the platform foresaw needing large amounts of storage. This poll provided an insightful snapshot of the current state of the AnVIL and demonstrated areas where the AnVIL Team can take specific steps to address barriers related to platform adoption and further support the existing and varied AnVIL Community. This work can be built upon through user interviews, community discussion, and coordinating a recurring poll.

## Introduction

The exponential growth in biomedical data generation has created unprecedented challenges in data storage, analysis, and sharing [1,2]. Despite its potential for research and teaching applications [3], cloud computing platforms in biomedicine have historically faced adoption challenges. Would-be users struggle with technical complexity around data transfer and user interface [4], familiarity and onboarding [3], cost management [5], and data security concerns [4,6]. Understanding the user’s activity, their backgrounds, and perspective can help address these concerns and guide platform development, adoption, and impact [7–12]. For example, usage metrics collected by the Cancer Genomics Cloud have revealed time savings available to researchers in cloud-based genomic analysis environments [13]. Direct polling of users can also inform potential areas of improvement. The Galaxy Project team routinely gauges satisfaction with their hands-on activities [14][15]. Similarly, Nextflow community leaders have surveyed users to understand their transition to cloud computing and other preferences [16]. However, less is publicly known about users and their preferences on other platforms.

The NHGRI Genomic Data Science Analysis, Visualization, and Informatics Lab-space (AnVIL) [17] has emerged as a response to data storage, analysis, and sharing challenges, offering a federated cloud platform that integrates data management, analysis capabilities, and access controls [1]. While AnVIL and other cloud platforms are enabling the shift from traditional institutional computing clusters to cloud-based genomic analysis [18], understanding user experiences and potential challenges will be central to AnVIL’s ability to meet the needs of cutting-edge research [19,20]. Little is formally known about the types of users that leverage AnVIL, including their experiences, preferences, and awareness of documentation and resources.

As a comprehensive environment that includes Terra, Galaxy, RStudio/Bioconductor, Dockstore, workflows and Jupyter Notebooks, AnVIL offers multiple paths for data analysis and collaboration. However, flexibility also introduces complexity for the researcher choosing their analytical approach. AnVIL’s highly customizable environment also makes it challenging for platform developers to identify specific challenges users are having that can be met with actionable improvements. For example, developers might not know whether users are unfamiliar with the AnVIL implementation of the tool (such as “Bioconductor on AnVIL”) or are simply unfamiliar with the tool more generally. AnVIL’s innovative approach to data sharing, where researchers “move to the data” rather than downloading datasets locally, leads to more unknowns, such as which datasets are most in demand and what kinds of data (genomic, metabolomic, etc.) developers should keep in mind. The most important features of the AnVIL platform might also differ between returning and potential new users.

To systematically investigate user perspectives, we conducted a community poll targeting returning and potential AnVIL users. Our study aimed to:

1. Examine the background and current work of users to develop appropriate personas
2. Understand barriers to platform adoption and user preferences for training and support
3. Assess researchers’ technological comfort with cloud-based genomic analysis tools
4. Identify computational and data analysis resource needs

By providing a detailed understanding of user experiences, this user poll seeks to inform future platform development, enhance user support strategies, and ultimately accelerate the adoption of cloud-based genomic analysis technologies.

## Methods

### Design of the poll

Given that AnVIL is a newer platform that has never polled users before, this community poll was designed to collect baseline information specifically with regards to the aforementioned study aims (Table 1). Questions were sourced from the AnVIL team as a whole – a combined effort across Johns Hopkins University, the Broad Institute, Fred Hutch Cancer Center, Vanderbilt University Medical Center, and other institutions within the AnVIL team.

**Table 1:** Relation of study aims to the design of the State of the AnVIL 2024 Community Poll. Sections of the State of the AnVIL 2024 Community Poll (Part 1, Part 2, etc.) are listed in the first column as the row names. Column names are the enumerated study names. X’s are added at the intersection of a poll section and a study aim if questions within that section are relevant to the study aim. <u>Supplemental material</u> provides a further breakdown of how each question relates to the study aims.

|  | Examine the background and current work of users to develop appropriate personas | Understand barriers to platform adoption and user preferences for training and support | Assess researchers' technological comfort with cloud-based genomic analysis tools | Identify computational and data analysis resource needs |
| --- | --- | --- | --- | --- |
| Part 1:<br>Self-identify | x |  |  |  |
| Part 2:<br>Feature importance +<br>Returning user specific Q's | x | x | x | x |
| Part 3:<br>Demographics | x |  |  |  |
| Part 4:<br>Experience | x |  | x | x |
| Part 5:<br>Awareness |  | x |  |  |
| Part 6:<br>Preferences |  | x | x | x |

The AnVIL Community Poll was constructed using Google Forms. The poll had 6 parts (Figure 1, Table 1). In the first part, using a multiple choice question, respondents were asked to select the best description of their current usage of the AnVIL platform. We used the answers to categorize respondents as returning or potential users of the AnVIL platform. Going forward, we discuss usertypes as either returning or potential users of the AnVIL. We define returning users as those who selected any of the following: “For ongoing projects (e.g., consistent project development and/or work)”, “For short-term projects (e.g., short, intense bursts separated by a few months)”, or “For completed/long-term projects (e.g., occasional updates/maintenance as needed)”; and potential users as those who selected any of the following: “I do not currently use the AnVIL, but have in the past”, “I have never used the AnVIL, but have heard of it”, or “I have never heard of the AnVIL”. Depending on the answer to this question, respondents were directed to the second part as either a returning or potential user of the AnVIL platform. Part 2 of the poll had a single question for potential users or 7 questions for returning users. The 6 additional questions asked about their experience with, needs, and recommendation likelihood for using the AnVIL while both categories of respondents were asked to rank features of the AnVIL according to their importance for their continued (returning user) or potential (potential user) use of the platform. Every respondent was given the same questions for the remaining parts of the poll (parts 3-6), with 5, 5, 2, and 6 questions respectively for each section. If respondents assented that they were willing to be contacted again to give input in the future, then they were directed to provide an email address. Finally, all users were provided with a link to a separate google form in case they wanted to be entered in a potential prize raffle. Due to there being a total of 20-26 questions, we estimated the poll would take 10-15 minutes to complete (∼30 seconds/question).

**Figure 1:**
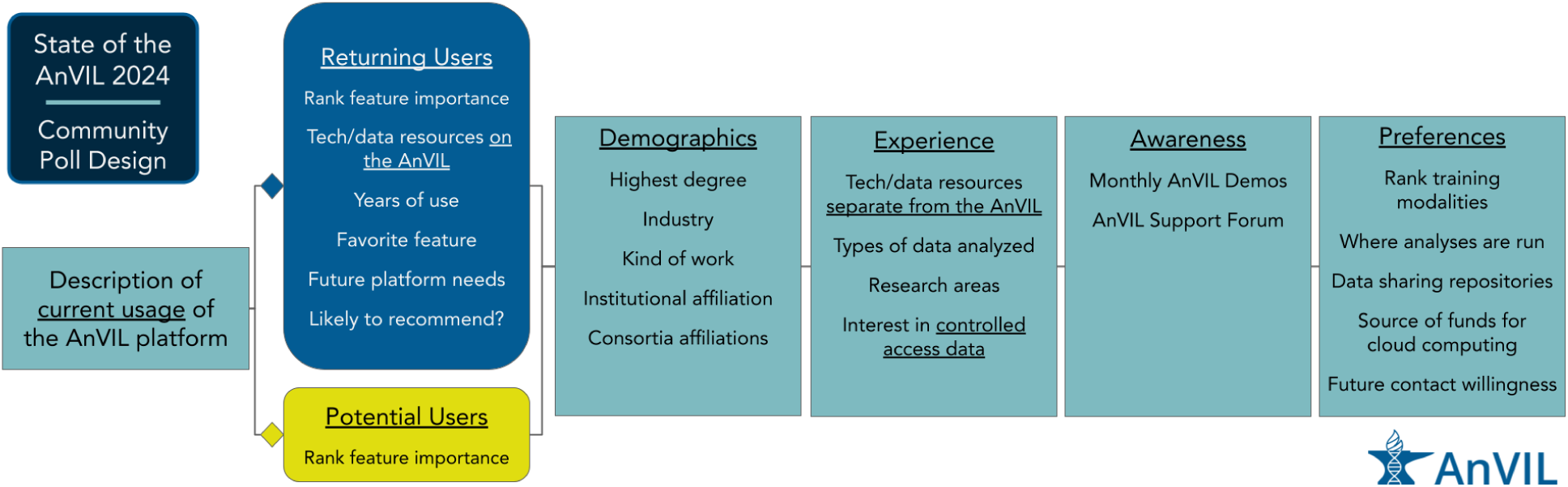
S**t**ate **of the AnVIL 2024 Community Poll Design.** The State of the AnVIL 2024 Community Poll was designed such that there were 6 parts. Every respondent was asked the same questions for Parts 1, 3, 4, 5, and 6 while respondents were provided with user type specific questions in Part 2, depending upon their answer to the first question of the poll. Part 1 was used to classify a respondent as a returning or potential user of the AnVIL; Part 2 asked questions specific to each user type (returning or potential); Parts 3 - 6 contained demographics, experience, awareness, and preferences questions respectively. Brief descriptions of questions within each part are provided.

**Figure 2:**
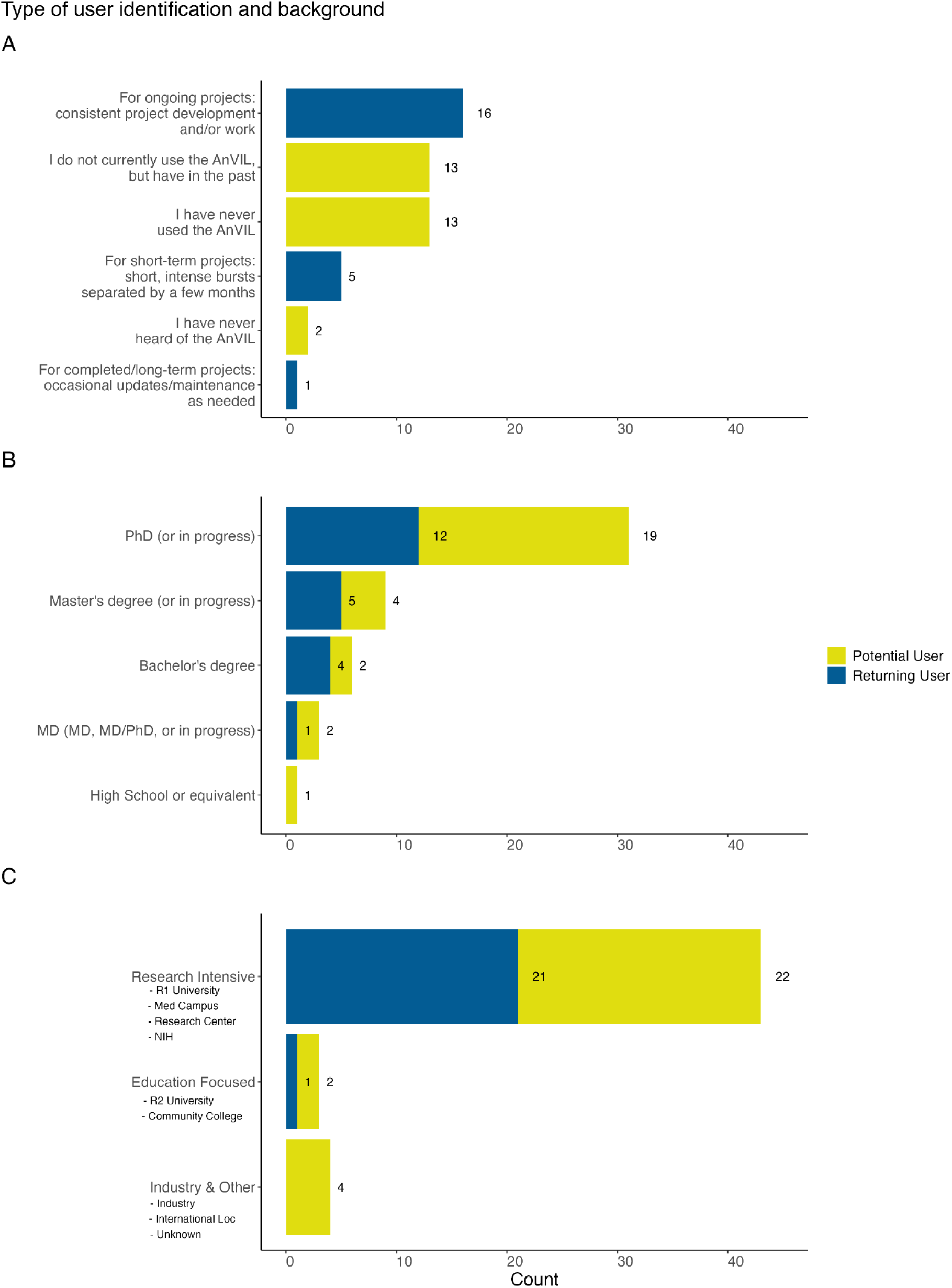
Background of users in our sample. A: We asked all respondents “How would you describe your current usage of the AnVIL platform?” Respondents were identified as returning or potential AnVIL users based on their responses. Most of the 22 returning users leverage AnVIL for ongoing projects. The 28 potential users were evenly split between those who have never used the AnVIL (but have heard of it) and those who have used the AnVIL previously, but don’t currently. B: We asked “What is the highest degree you have attained?” Most of the respondents have a PhD or are currently working on a PhD, though a range of career stages were represented. C: We asked all respondents “What institution are you affiliated with?” Most of the respondents, but also the majority of returning individuals using the AnVIL, reported being affiliated with a research intensive institution.

**Figure 3:**
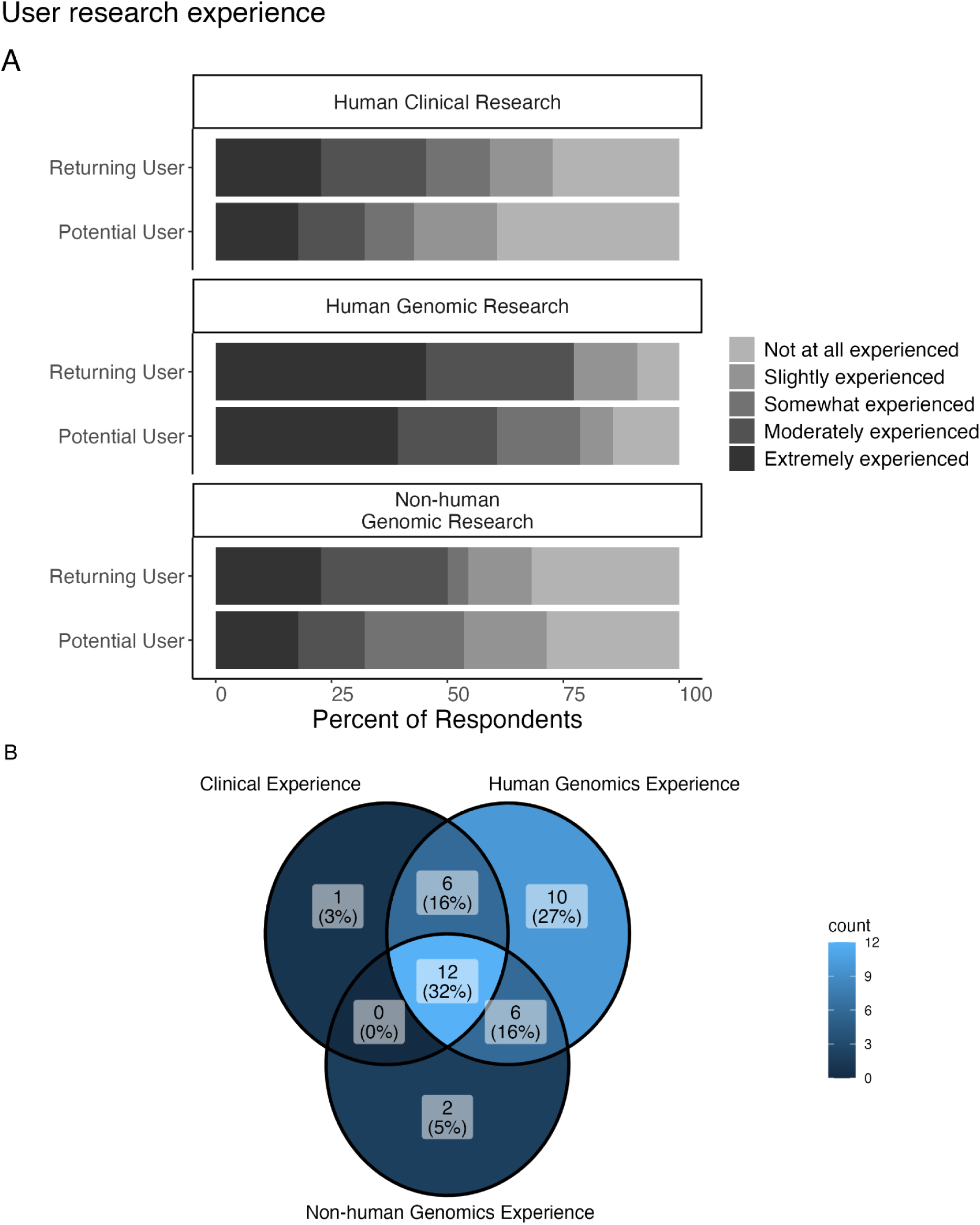
**Background of users and researcher’s technological comfort**. A: We asked respondents “How much experience do you have analyzing the following data categories?” 21 respondents report that they are “extremely experienced” in analyzing human genomic data, while only 6 respondents report that they are “not at all experienced” in analyzing human genomic data. However, more respondents report being “not at all experienced” in analyzing human clinical data and non-human genomic data. B: Venn diagram showing the overlap for respondents who reported being “moderately” or “extremely experienced” for these various research categories (n = 37). 32% reported such levels of experience for all 3 research categories; the next highest percentage (27%) reported such levels of experience for human genomic data only with no overlap with other research categories.

**Figure 4:**
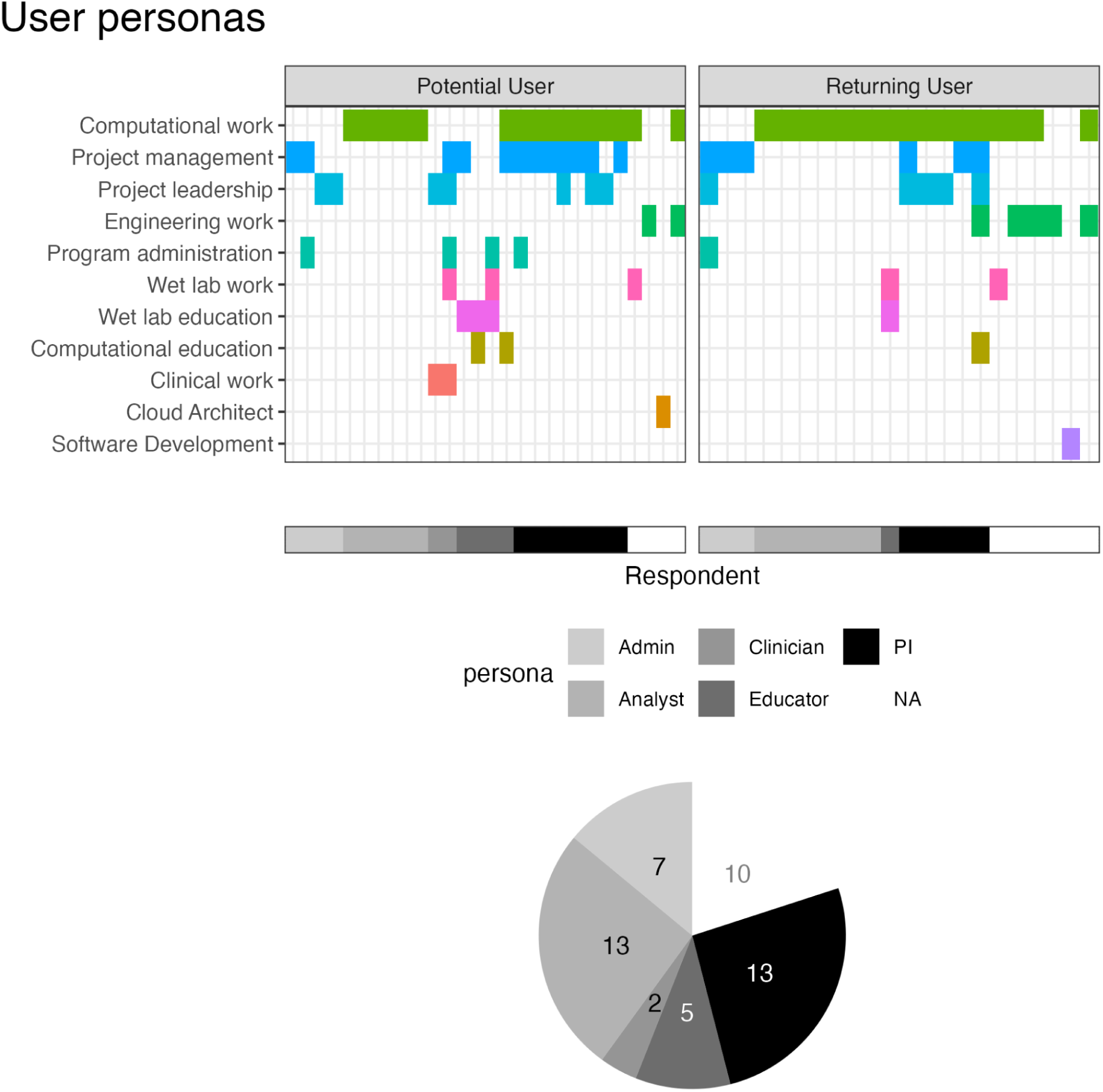
Current work of users related to appropriate personas. We asked all respondents “What kind of work do you do?” Possible selections (computational work, computational education, project management, etc.) are shown on the y-axis and cells are colored for each choice a respondent (x-axis) selected. Based on selections, respondents were clustered into Admin, Analyst, Clinician, Educator, PI, and not assigned (NA) personas. These assignments are shown in grayscale along the x-axis with a corresponding pie chart showing the relative abundance of these assignments. 2 potential users were assigned “Clinician” personas and 4 were assigned “Educator” personas, compared to 0 and 1 respectively for returning users. The other personas show similar abundances between potential and returning users.

The outline of the poll is as follows (Figure 1):

Part 1: Self-identify as a returning or potential user

Part 2: Rank existing or potential features according to their importance (everyone) + returning AnVIL user specific questions about experience with the platform

Part 3: Demographics (e.g., degree level, type of work, institution, consortia affiliations, etc.)

Part 4: Experience (e.g., general bioinformatics experience, relevant datasets, etc.) Part 5: Awareness and utilization of available AnVIL support

Part 6: Preferences (e.g., training modality, analyses platforms, etc.) For a pdf version of the full poll, see this <u>supplementary file</u>.

### Recruitment for/Dissemination of the poll

The AnVIL Community Poll was disseminated through multiple channels including posting on social media platforms like X (formerly Twitter) and LinkedIn, emailing applicable mailing lists (registered AnVIL users through Terra, AnVIL Demo attendees, AnVIL mailing list, etc.), and posting within Slack workspaces (AnVIL, nf-co.re, Community Bioconductor, Fred Hutch Cancer Center, and the Johns Hopkins University Genomics and Biostat communities). We also posted an advertisement for the survey as a news item on the anvilproject.org portal and the AnVIL Support Forum (help.anvilproject.org). Finally, the poll was sent directly to some select individuals and consortia who were asked to further circulate it. All advertisements mentioned the possibility of a prize raffle. The poll was conducted in Spring 2024. It was administered using Google Forms and open for responses from February 15th to March 25th.

Because of the nature of dissemination (through social media), we cannot confidently calculate a response rate. However, we consider this to be a data gathering exercise where we can report back the findings to the AnVIL Community and see if and how they resonate.

### Ethical Considerations

This work has been reviewed by The Johns Hopkins University Homewood IRB and was determined to not-qualify as human subject research (#HIRB00020632).

### Analysis of the poll

Following the closing of the poll, the Google sheet with form responses was locked so that edits could not be made and the results were imported from there for analysis using R. Respondent email (if provided) was used to de-duplicate responses. Identifying information or information that was highly specific and potentially identifiable like specific institution names was removed (email addresses) or annotated and grouped into categories (type of institution, e.g., an R1 University, R2 University, Community College, etc.) before removal.

Respondents were asked about their level of experience with human genomic, non-human genomic, and clinical research. Possible responses included “Not at all experienced”, “Slightly experienced”, “Somewhat experienced”, “Moderately experienced”, and “Extremely experienced” (a 5 point Likert scale). If a respondent selected “Moderately” or “Extremely” experienced for any of these categories, they were assigned as someone with experience in that research category. This method of assigning experience is an example of a Top 2 Box simplification [21].

Poll takers were also asked to select the type(s) of work they perform regularly Possible choices for type(s) of work were Computational work, Engineering work, Wet lab work, Clinical work, Computational education, Wet lab education, Project leadership, Project management, Program administration, or Other (with free text entry if Other was selected). For analysis purposes, we grouped users into 5 possible personas: Clinician, Analyst, Educator, Leadership (PI), or Admin. These groups were created based on participant answers to the “type of work” question. A clinician persona was assigned if “Clinical work” was selected; An analyst persona was assigned if only “Computational work” was selected; An educator persona was assigned if “education” appeared in a response; A leadership persona was assigned if project leadership, management, or administration co-occurred with “Computational work”; and an Admin persona was assigned if any of project leadership, management or administration was selected without any other kinds of work selected. With this assignment procedure, some responses were not assigned personas and two were assigned multiple personas. To simplify those assigned multiple personas, the selections were individually inspected and categorized. We also performed principal components analysis, and did not observe any conspicuous clustering of respondents (see https://github.com/fhdsl/SOTA2024_ReportOut).

When respondents were asked to rank preferences or level of importance (Figure 5), responses were translated into rank values. If n choices were given, “Most preferred/important” was assigned a value of 1, with each subsequent value denoting decreasing preference or importance. The “Least important/preferred” was assigned the largest possible value. An overall or average rank choice was assigned by summing the rank values and dividing by the total number of respondents. For an average knowledge or comfort score (Figure 7a), a similar procedure was followed; however, for these questions the higher value was associated with greater knowledge or comfort and a score of 0 was used for responses of not knowing at all. Rank and comfort scores were averaged within user cohorts (e.g., persona or user type) rather than considering all respondents together.

**Figure 5.**
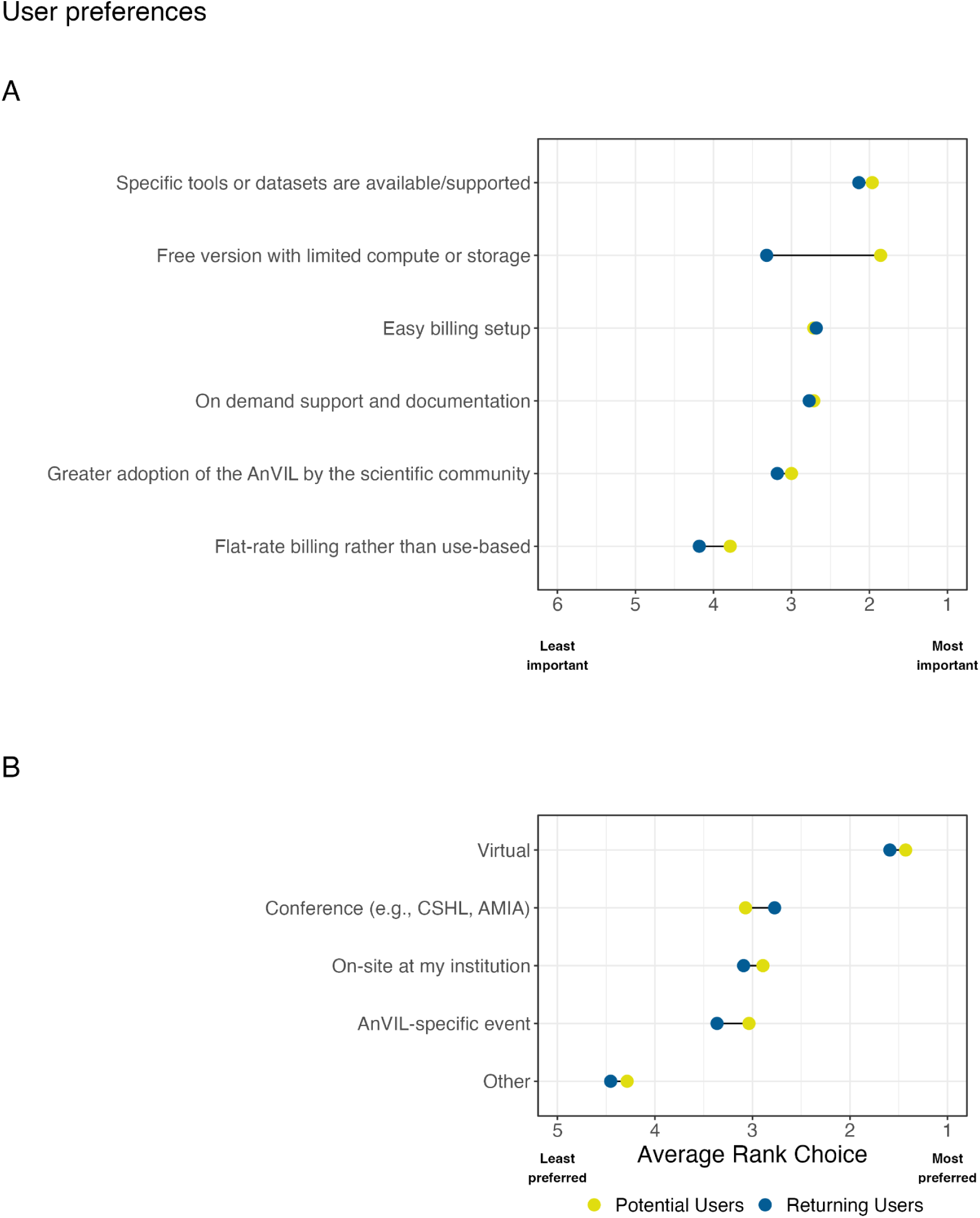
Barriers to platform adoption and user preferences for training and support in our sample. A: We asked all users to “Rank the following features according to their importance to you as a potential user or for your continued use of the AnVIL.” Responses were averaged within potential and returning user cohorts to find an average rank. All respondents rated having specific tools or datasets supported/available as an important feature for using AnVIL. Compared to returning users, potential users rated having a free-version with limited compute or storage as the most important feature for their potential use of the AnVIL. B: We asked all respondents “Rank how/where you would prefer to attend AnVIL training workshops.” Responses were averaged within potential and returning user cohorts to find an average rank. Both returning and potential users preferred virtual training workshops over other modalities.

**Figure 6:**
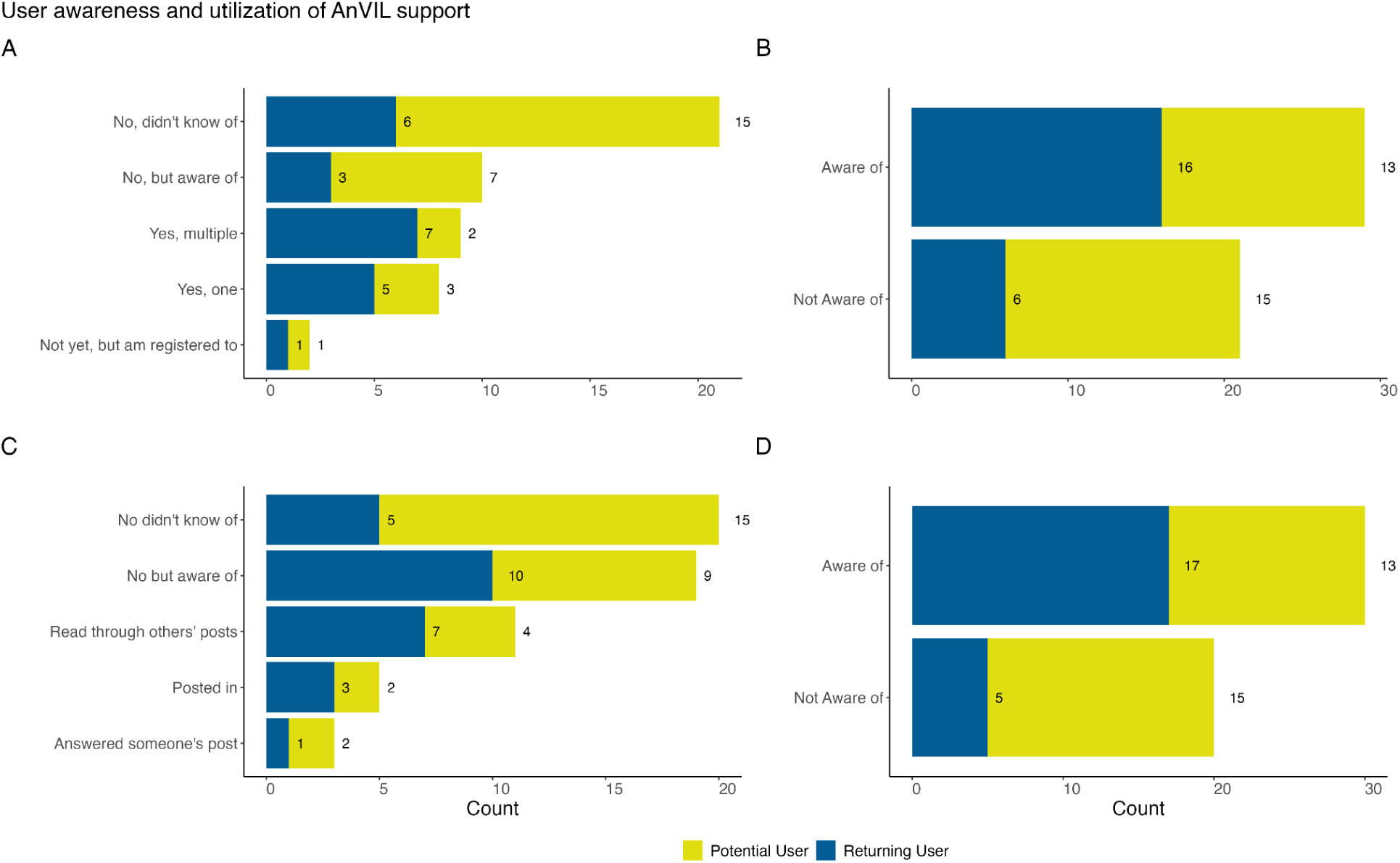
Awareness and utilization of training and support. A & B: We asked all respondents “Have you attended a monthly AnVIL Demo?” A: Most respondents had not attended an AnVIL Demo. However, returning users were more represented among AnVIL Demo attendees. B: All responses to (A) except “No, did not know of” were aggregated, showing that the majority of respondents are aware of AnVIL Demos. C & D: We asked all respondents “Have you ever read or posted in our AnVIL Support Forum?” C: Raw responses are shown (users could select more than one). Most respondents have not used the AnVIL support forum, but utilization in some form is reported by 24% of respondents; reading through others’ posts is the most common way of utilizing the support forum within this sample. D: Each set of user responses are recoded and aggregated to examine whether users are or are not aware of the AnVIL Support Forum. We observe that there is awareness of the support forum across potential and returning users.

**Figure 7:**
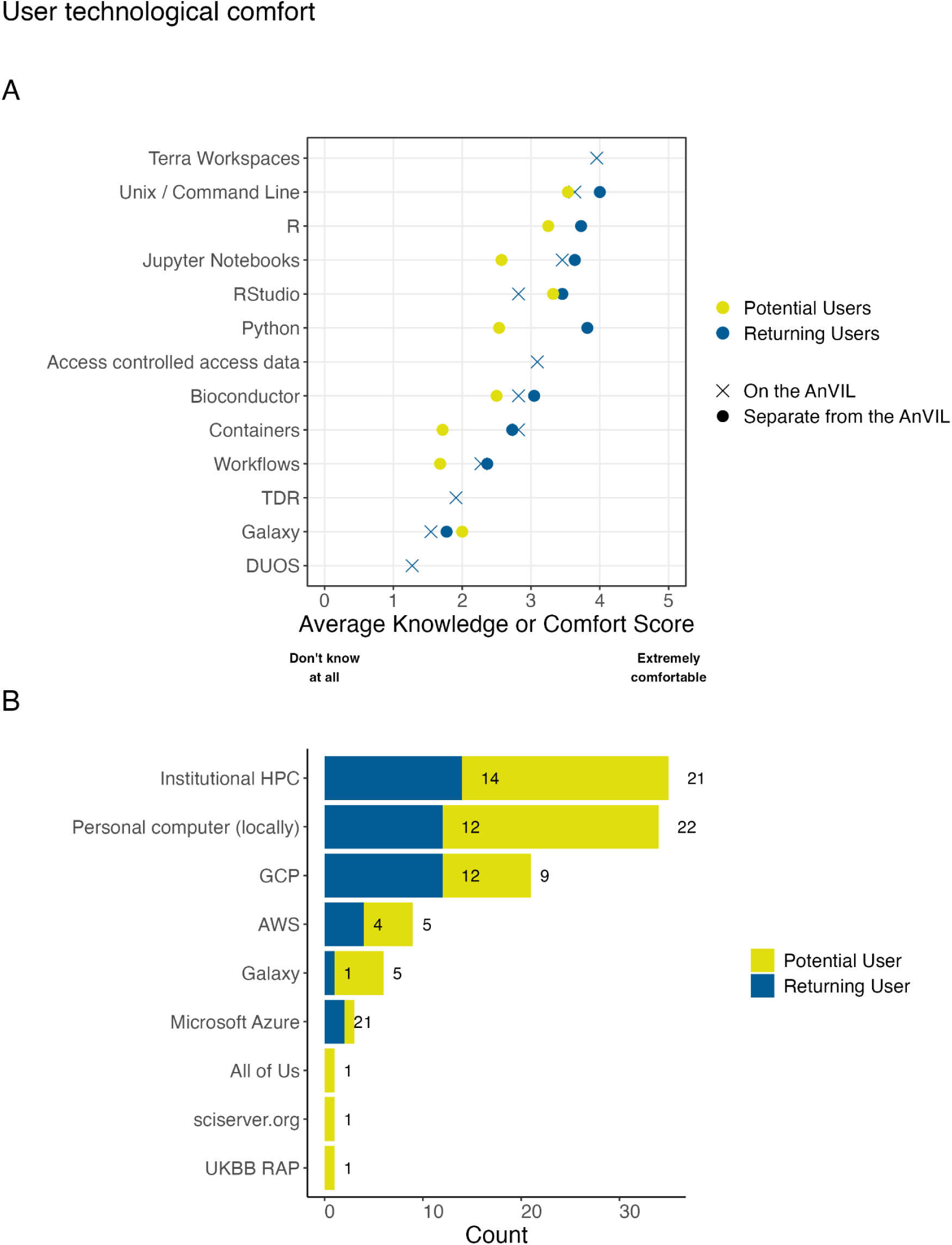
Technological comfort with cloud-based genomic analysis tools. A: We asked respondents “How would you rate your knowledge of or comfort with these technologies or data features?” Except for Galaxy, potential users tended to report lower comfort levels for the various tools and technologies when compared to returning users. Overall, there was less comfort with containers or workflows than using various programming languages and integrated development environments (IDEs). B: We asked all respondents “Where do you currently run analyses?” Institutional HPC and locally run (personal computers) were the most common responses. Google Cloud Platform (GCP) was reported as used more than other cloud providers within this sample. We also saw that potential users reported using Galaxy (a free option) more than returning users do.

In Part 5 of the poll, respondents were asked about their experiences with user support options. Respondents could select “No, didn’t know of”, “No, but aware of”, or some specific description related to how they used the support option (e.g., attending a demo or answering someone’s post on the support forum). We broadly translated these responses into two binary reporters: one for awareness and the other for active use. To understand awareness, we combined responses in the following way: Anything other than “No, didn’t know of” was combined to represent awareness. To understand use, we combined responses such that anything other than “No, didn’t know of” and “No, but aware of” were examples of use or future use. If looking at past use (ignoring future use), “Not yet, but registered to” was also not included in answers representing use. (See the <u>supplementary material for a table</u> representing these categorizations.)

*All code for reproducing the stats and figures in this analysis are available on GitHub:* https://github.com/fhdsl/SOTA2024_ReportOut*. A de-identified dataset is also available*.

## Results

### Responses

This AnVIL user poll was conducted in Spring 2024, receiving 52 total responses. Two responses were determined to be submitted by duplicate users, leaving a total of 50 user responses used in the analysis. Given how the AnVIL Community Poll was advertised through general, often public channels rather than enumerated closed lists, it was not possible to compute an exact response rate. Considering the distribution of daily responses compared to recruitment efforts for the poll, responses were received on days without advertisement, but days with advertisement (at least 7 unique days) aligned with or directly preceded receiving responses. The two days with the highest response count (7 and 8 responses) aligned with a user listserv reminder email (2024-March-13) and an X (formerly Twitter) post (2024-March-14). The average number of responses in a day was about 1, though the most common number of responses was 0 (21 different days), followed by 1 (8 different days) if restricted to non-zero responses. When considering the institutional affiliation of respondents, there were no higher than 2 responses from users from the same institution on any day. Both potential and returning users responded throughout the time period with no discernible pattern related to recruitment method.

### User backgrounds and current work

Of the 50 de-duplicated responses, 44% (n=22) were from returning users and 56% (n=28) were from potential users. The majority of returning user responses (73%, n=16) belonged to the group who use the AnVIL for ongoing projects, with consistent work on the platform. The majority of potential users were evenly split between two groups (46%, n=13 each), those who have never used the AnVIL (but have heard of it) and those who have used the AnVIL previously, but don’t currently (Figure 2A).

Most respondents in our sample had obtained or were in the process of obtaining a PhD (62%, n=31) though a range of career stages were represented (Figure 2B) including those with little to no post-secondary education and those with multiple advanced degrees. Clinical advanced degrees (MDs) (6%, n=3) were less represented than either research degrees (PhD) or Master’s degrees (18%, n=9).

Most of the respondents using the AnVIL (n=21) reported being affiliated with a research intensive institution (e.g., an R1 University, a medical campus, a research center, or the NIH), while one reported being affiliated with an education focused institution (e.g., R2 University or community college). Only potential users of the platform reported being affiliated with an industry-based institution (Figure 2C). In our sample, we observed very few industry partners.

Of the 50 responses, 21 provided consortia affiliations across 23 unique affiliations (respondents could report more than one consortium). Of the 21 responses providing consortia affiliations, 13 were from returning users. Genomics Research to Elucidate the Genetics of Rare Diseases (GREGoR)[22,23], Polygenic Risk Methods in Diverse Populations (PRIMED)[24,25], Electronic Medical Records and Genomics (eMERGE)[26], Centers for Common Disease Genomics (CCDG)[27] and The Genotype-Tissue Expression Project (GTEx)[28] were the top 5 most represented with n=3 each for GREGoR, PRIMED, and eMERGE and n=2 each for CCDG and GTEx. Additionally, all 5 of these consortia were represented in the subset of consortia affiliations reported by returning users.

When asked about experience with analyzing human genomic, human clinical, and non-human genomic data, 21 respondents report that they are extremely experienced in analyzing human genomic data, while only 6 respondents report that they are not at all experienced in analyzing human genomic data. However, for human clinical data and non-human genomic data, more respondents report being not at all experienced in analyzing those data than report being extremely experienced (Figure 3A). Returning and potential users showed a similar distribution on experience levels across these 3 research categories. If we consider the Top 2 Box simplification, lumping together “Moderately” and “Extremely” experienced responses (the highest 2 of the possible rankings), returning users show slightly more experience across all 3 research categories.

When limiting the responses we consider to include only respondents who reported being “moderately” or “extremely” experienced for at least one of the categories, we are left with 37 responses (17 from returning users). Considering the overlap among experience within these categories for these responses, 32% (n=12) reported these levels of experience for all 3 research categories and 27% (n=10) reported this level of experience for only human genomic research (but no other research categories) (Figure 3B). 16% (n=6) reported this level of experience for both human clinical and human genomic research (but not non-human genomic research) and another 16% (n=6) reported this level of experience in analyzing both non-human and human genomic research.

When asked to select the kind of work that they performed, 44% (n=22) selected only 1 description, 36% (n=18) selected 2 descriptions, 16% (n=8) selected 3 descriptions, and 4% (n=2) selected 5 descriptions. “Computational work” was the most frequently selected response, reported by 68% of respondents (n=34). “Project management” and “Project leadership” followed at 36% (n=18) and 24% (n=12) respectively.. Of the 22 respondents who selected only 1 description, “Computational work” was the most frequently selected (n=13, 59%). Of the responses who selected more than one description, “Computational work” was paired most frequently with “Project management” or “Project leadership”. (n=11 and 7 respectively). Two respondents utilized the “Other” option, each supplying one work description. These included “Cloud Architect” and “Software Development”. The distribution of work descriptions selected were similar when comparing between potential and returning users (Figure 4).

All of the results summarized so far describe the varied backgrounds and current work of respondents in our sample. Despite this variety, we expect several overarching user groups: Admins, Analysts, Clinicians, PIs, and Educators. Given that the kind of work question focused on work activities and that these expected user groups have differential work activities, we assigned personas based on the responses to the kind of work question (Figure 4). 10 respondents were left uncategorized, but the majority of respondents were assigned to either the Leadership (PI) persona (n=13, showing evidence of computational work together with some form of project management, leadership, or administration) or the Analyst persona (n=13, only reporting computational work). 7 respondents were categorized as an Admin persona. The Leadership (PI), Analyst, and Admin personas showed a similar split of assignment across potential and returning users. The Clinician and Educator personas were differentially represented in this sample: 2 potential users were assigned a Clinician persona because of selecting clinical work (this option was not selected by returning users) and 4 potential users were assigned an Educator persona while only 1 returning user was.

### Barriers & user preferences

The kind of work that researchers perform or different hats that respondents wear are likely to influence their preferences and barriers to adoption. Respondents were asked about features that are most important for their continued or potential use of the AnVIL. Of the features listed, currently the AnVIL utilizes use-based billing and does not offer a free version. When looking at responses specific to the assigned personas, we observe that the average rank choice for easy billing setup is highest for Admins compared to other personas; Educators and Clinicians rank both a free version with limited compute or storage and greater adoption by the scientific community more highly than other personas; and PIs rank having specific tools or datasets available/supported as their most important feature of AnVIL. Specific tools or datasets being available and supported ranks as the most important feature for both returning and potential users of the AnVIL (Figure 5A). Potential users rank having a free version of the AnVIL with limited compute or storage more highly than returning users do. Easy billing setup and on demand support and documentation are ranked in the middle with similar preferences observed across user types. Flat-rate billing rather than use-based billing is ranked the lowest across this set of features within this sample.

As for respondent preferences regarding training modalities/locations, we see that the vast majority of responses in our sample rank virtual options above all other modalities (Figure 5B).

Respondents viewed conferences, on-site at their institution, or AnVIL specific events with similar preference ranks. These observations were consistent across user types. At the time of this poll, the AnVIL had offered training through every represented modality option except for an AnVIL-specific event (e..g, the AnVIL Community Conference whose inaugural event occurred after the poll closed).

The AnVIL already supports several virtual training opportunities including monthly AnVIL Demos and a 24/7 online support forum (help.anvilproject.org). The monthly AnVIL Demos are virtual meetings hosted over zoom where typically the first 30 minutes are used for a demonstration highlighting what is possible on the AnVIL and the last 30 minutes are reserved for questions and community discussion. The AnVIL Demos are advertised through the <u>AnVIL</u> <u>mailing list</u>, on the <u>events tab of the AnVIL portal</u>, and <u>described on the support forum</u>.

Recordings are posted to YouTube and listed as part of the <u>AnVIL Collection</u>. The 24/7 online support forum provides a place for users to submit questions or read through/reply to threads of questions regarding the platform. The AnVIL outreach team responds to questions and sorts threads into relevant categories (e.g., data access, feature requests, etc.). When asked about their utilization of these virtual training opportunities, we found that more respondents were aware of these offerings than were not aware. Potential users were split at 46% (n=13) knowing about monthly AnVIL Demos or the support forum and 54% (n=15) not being aware of the support opportunities (Figure 6B,D). Interestingly, only 71% (n=20) of potential users had matching aware/not aware responses for both AnVIL Demos and the support forum. Of the 15 not aware respondents, only 73% (n=11) were not aware of both AnVIL Demos and the support forum. Similarly, 86% (n=19) of returning users had matching aware/not aware responses for both AnVIL Demos and the support forum. Only 4 returning user responses were not aware of both the AnVIL Demos or the support forum. Returning users showed a higher percentage of respondents who were aware of the support opportunities (n=16, 73% and n=17, 77%) than potential users (Figure 6B,D). Returning users are also more represented in the group of respondents who have attended AnVIL Demos (n=12 or 71% of attenders)(Figure 6A) or utilized the support forum (n=8 or 67% of those who have read through, posted, or replied) in this sample. In general, most poll respondents have not attended AnVIL Demos with only 34% (n=17) reporting attendance (Figure 6A). For those who have attended an AnVIL Demo within this sample, attending more than one demo or just one demo occurs at similar rates (Figure 6A). Most respondents have not utilized the support forum with only 24% (n=12) in some way reporting use. Reading through others’ posts is the most common form of utilization (Figure 6C).

### User technological comfort

The poll asked about comfort with technology in two different sections: within Part 2 (the returning user exclusive questions) and within Part 4 (Experience). Within Part 4, all respondents were asked to rank their knowledge or comfort with certain technologies separate from the AnVIL. Only respondents identified as returning users were asked about these same technologies (excluding specific programming languages) on the AnVIL. Returning AnVIL users were the only respondents asked about knowledge of or comfort with AnVIL specific features such as Data Use Oversight System (DUOS), Terra Data Repository (TDS), Terra Workspaces, or generally accessing controlled access data. Returning users reported higher comfort levels for the various tools and technologies when compared to potential users. The one deviation from this trend is a slight increase in the comfort with Galaxy reported by potential users.

Overall, there is less comfort with containers or workflows than various programming languages and integrated development environments (IDEs). While respondents report fairly high comfort with Jupyter Notebooks and RStudio for interactive analysis on the AnVIL, Galaxy on AnVIL is among the technologies with the lowest average knowledge or comfort. Returning users report low levels of knowledge or comfort with TDR and DUOS as well. They report intermediate levels of comfort with accessing controlled access data in general and the highest level of comfort with workspaces.

All respondents in our sample were asked where they currently run analyses. Respondents could select multiple choices. Institutional High Performance Computers (HPCs) (70%, n=35; 64% of returning users and 75% of potential users) and locally on personal computers (68%, n=34; 55% of returning users and 79% of potential users) were reported at the highest rates in this sample. For provided cloud options, Google Cloud Platform (GCP)[29] was reported as used the most (n=21) split across user types (55% of returning users and 32% of potential users) of the AnVIL, compared to 22 returning AnVIL users in our sample). We also observed that potential users report using Galaxy (a free option)[30](18%, n=5), more than returning users do (5%, n=1). Potential users also submitted All of Us [31], sciserver.org, and UKBB RAP [32] as other platforms they use to perform analyses.

### *User* resource and data needs

Returning users (n = 22) were asked about their foreseeable computational needs. The most common response was needing large amounts of storage (e.g., Terabytes), selected by 50% (n=11); 32% (n=7) only selected this option. Many nodes, GPUs, and large memory (>192 GB RAM) were also reported as foreseeable needs, but to a lesser extent (selected by 27%, 27%, and 18% respectively). Only 1 response selected all 4 computational needs. The most common combination of selections was GPUs and many nodes (n=2). All other combinations were only selected once or not at all (though this could be an artifact of a small sample size and 11 possible combinations of multiple options).

46% (n=23) of all respondents flagged the AnVIL as a repository that they use or are considering use of to share data to comply with the NIH DMS policy. Of those 23, 65% (n=15) were returning users of the AnVIL (68% of the returning user sample of this community poll).

Over half of all respondents (n=29) reported that they are “extremely interested” in working with controlled access datasets. When provided choices of specific controlled access datasets that respondents have accessed or are particularly interested in accessing with AnVIL, All of Us [33] and UK Biobank [34] were the most selected (n=34 each). GTEx was selected by 64% (n=32) of respondents and CCDG was selected by 40% (n=20). All of Us, UK Biobank, and GTEx remained the top three when we subset the data to consider (1) only responses from those moderately or extremely experienced with human clinical data, (2) only responses from those moderately or extremely experienced with human genomic data, and (3) only responses from those moderately or extremely experienced with non-human genomic data. CCDG remained the 4th rank selection for those moderately or extremely experienced with non-human genomic data, but was surpassed by eMERGE and HPRC [35] when considering the subsets of researchers experienced with human clinical and human genomic data. Note that poll respondents were informed that All of Us and UK Biobank were not currently available on the AnVIL due to policy restrictions. (Note: Since the results of this poll were analyzed, All of Us is now more accessible to researchers. [36])

When asked about interest in different types of data, respondents selected Genomes/exomes (88%, n=44) and Transcriptomes (62%, n=31) as the major types of data they would be interested in analyzing with the AnVIL. A variety of other options (e.g., Phenotypic, Single Cell, Electronic Health Record, Epigenomes, Metabolomes, Proteomes, Imaging, etc.) were selected at a rate of at least 10% (n=5), but no more than 40% (n=20). Survey, structural, and variant calling were the only options selected or mentioned by fewer than 10% of respondents.

## Discussion

These results together show that while the broad majority of AnVIL users in our sample share commonality in their research interests and needs from the AnVIL platform (e.g., high storage volumes, use as a data repository, ability to access controlled access datasets, etc.), the AnVIL Community maintains a wide range of expertise and research interests. The study found opportunities for platform adoption as well as areas where training should be enhanced to better support the growing community (Table 2, Figure 8).

**Figure 8:**
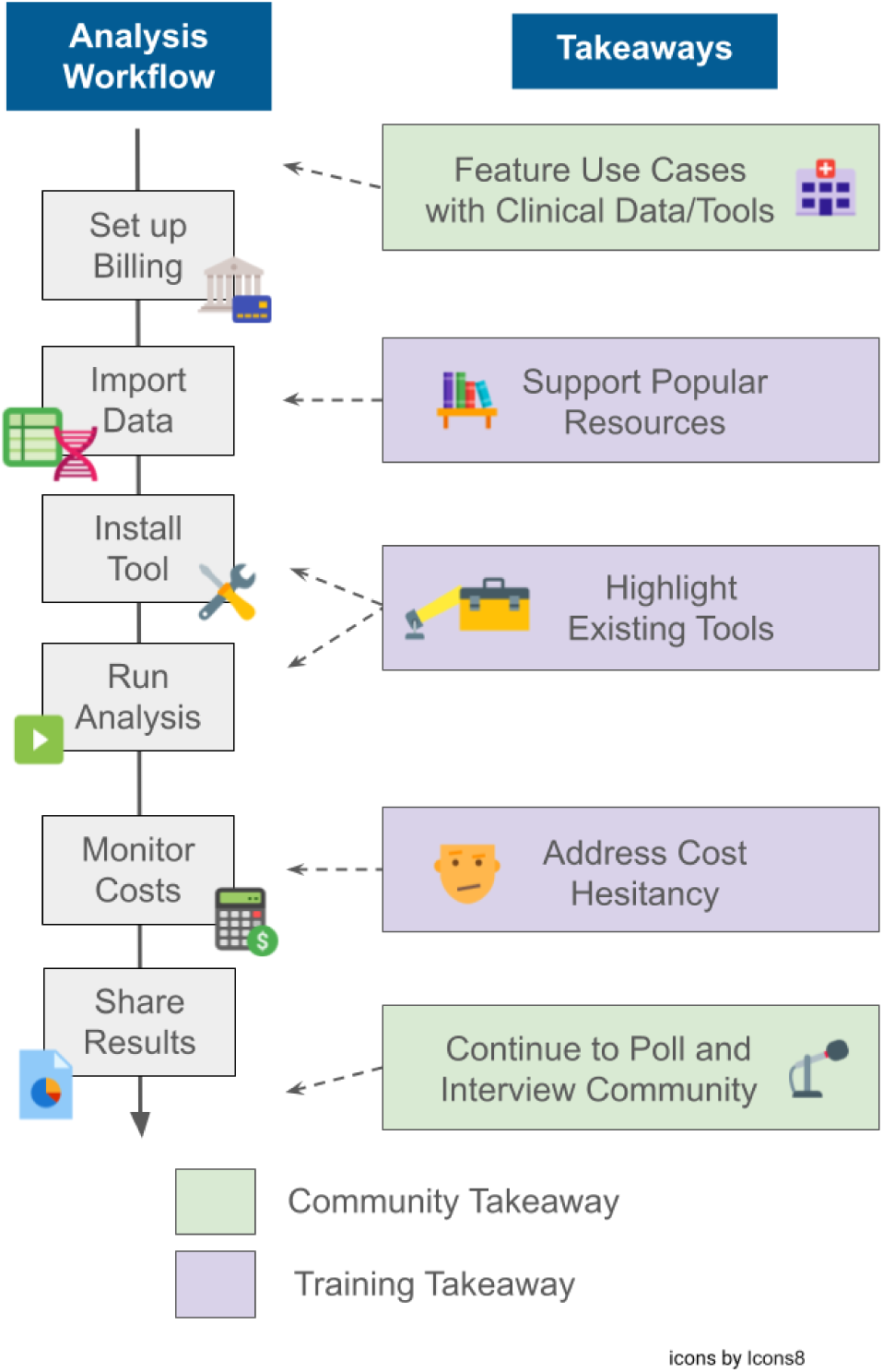
Relating poll takeaways to steps in a typical analysis workflow using the AnVIL platform. Poll takeaways from Table 2 are condensed to combine the takeaway with the proposed step(s) to address it and categorized as training or community takeaways. Training takeaways relate to those with steps the AnVIL Team can take to create new and enhance or highlight existing training materials to address. Community takeaways relate to those meant to converse with, learn from, or grow the AnVIL Community. These takeaways are aligned along a typical analysis workflow that may be performed on the AnVIL platform.

**Table 2:**
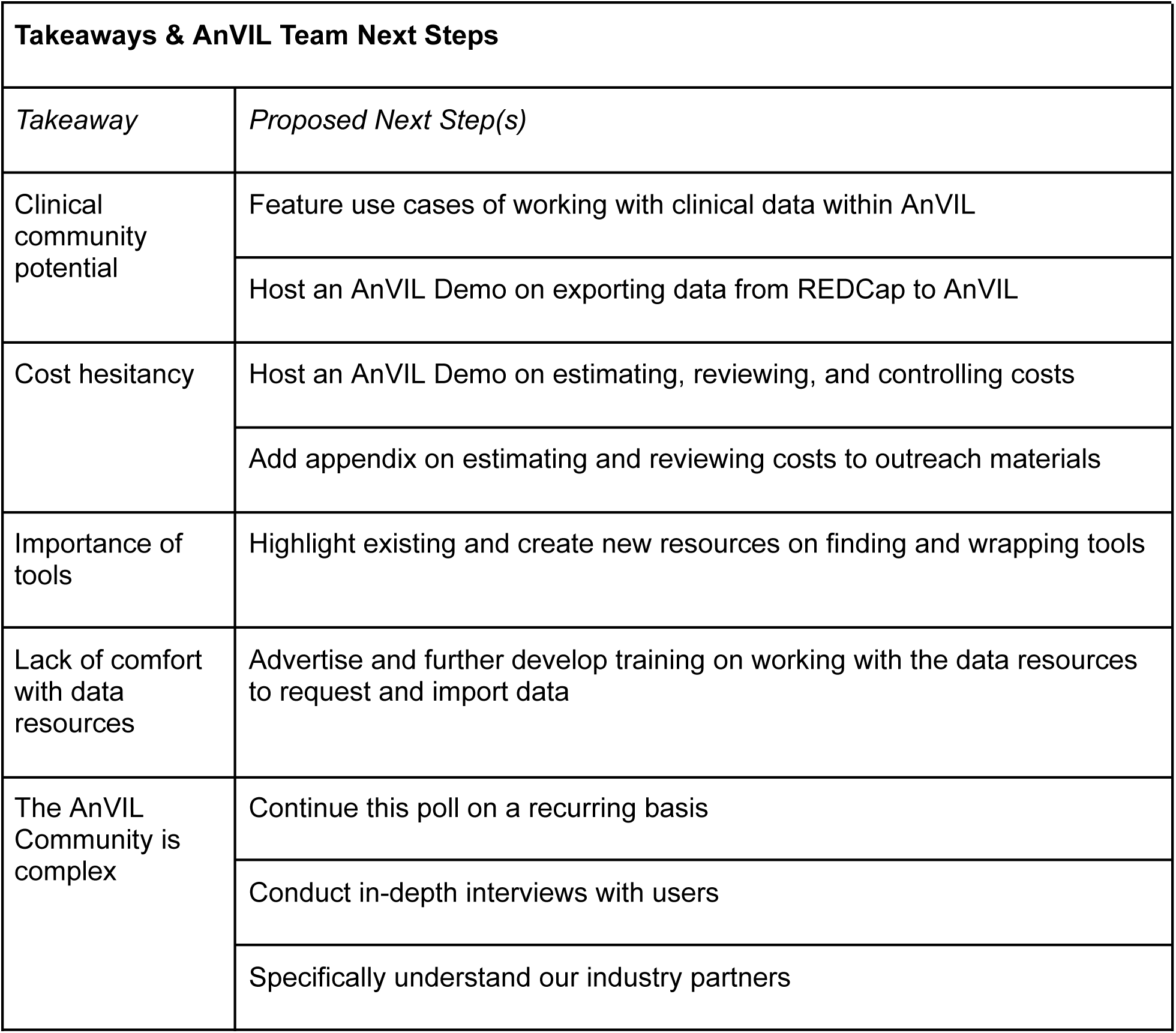
Takeaways and next steps for the AnVIL Team. Takeaways from analysis of the poll responses are listed in the first column as the row names. The second column proposes specific examples of next steps that the AnVIL Team can take to address these points.

Our first aim with this poll was to examine the background and current work of users. Given that AnVIL was developed to integrate and analyze large human genomics datasets and other biomedical data [17], we would expect that users have backgrounds and current interests aligned with human genomics data and research. In general there is a close alignment between what we observe and this expectation. Returning users in our sample did show slightly more experience with human genomic research when compared to non-human genomic and clinical research categories. This could be because AnVIL was designed in partnership with NHGRI, which focuses on human genomics, so respondents are likely to have more experience in that area generally. We also observed that returning users felt more experienced in research overall, which could be a potential source of response bias.

In addition to supporting human genomics research, there is also an interest in the AnVIL Community to increase the usage of the platform by clinicians and clinical researchers for biomedical data and analysis in general. Expanding the AnVIL Community to include clinicians could facilitate interdisciplinary breakthroughs in fields like precision medicine [37–39]. While our sample seems to be predominantly represented by human genomics researchers, we observe some respondents (both returning and potential users) with experience or interest in clinical data and research. For example, we observed (1) respondents with professional degrees, (2) those who selected clinical work as a kind of work they perform, (3) the representation of those who report they are moderately or extremely experienced with human clinical data analysis, (4) the increase in interest among that clinical experience cohort for the clinical dataset Electronic and MEdical Records and Genomics Project (eMERGE), (5) eMERGE consortia affiliations, (6) Electronic Health Record and phenotypic data being data types respondents were moderately interested in analyzing with the AnVIL. This suggests an outreach opportunity, where the AnVIL team could feature use cases of working with clinical data within AnVIL or host a demo of exporting data from REDCap (a clinical data capture and management platform) [40], [41] to AnVIL. In addition, the AnVIL Team could partner with relevant consortia to try to train and bring in more of the clinical research community and continue development of features useful to the clinical community like the recent REDCap integration [42].

Moreover, we aimed to consider backgrounds and research interests in order to develop broad personas for users of the AnVIL. Given the small sample size of the poll, utilizing multiple questions to categorize or cluster respondents was infeasible and we opted for a simplified approach of parsing the responses to the question about kinds of work respondents performed. Personas help the AnVIL Team understand the projects that people are doing and therefore ways in which they may want to utilize the platform. Different types of research activities require different resources and features. In addition, personas and roles that researchers may be identified with are fluid. For example, a specific user may use AnVIL for a research project but also use the platform in order to run a workshop or as a learning environment supporting a class project. Therefore, we utilize personas within this analysis to confirm user groups that we would expect to see and identify potential areas where the AnVIL Team can prioritize outreach, conversations, or feature development. We were limited both by the small sample size and confirmation bias given the approach taken here [43,44]; therefore we recommend working closely with the community to listen to and understand their roles and needs.

Our second aim with this poll was to identify barriers to platform adoption and user preferences for training and support. Cost seems to be a barrier to platform adoption in our sample as evidenced by potential users ranking offering a free tier as an important feature. The AnVIL provides computing and storage billing in line with the amount of use rather than charging a flat rate for access. No free tier with limited compute or storage is offered. It is possible that potential users are unaware of the billing logistics utilized by the AnVIL platform while returning users would not have selected this feature as important knowing that it is not offered. The AnVIL is working to improve cost transparency and has provided documentation on controlling costs [45] as well as discussions on estimating the costs for interactive analyses [46] and a resource benchmarking how long common workflows take with specific datasets and the associated costs [47]. In addition, the AnVIL is developing a tool to help researchers estimate costs of a workflow [48]. The desire for a free tier might also reflect frustrations with administrative overhead associated with paid platforms, as “easy billing setup” was also ranked highly for returning and potential users.

Both returning and potential users ranked specific tools and datasets being supported as very important for using the AnVIL. AnVIL already supports a variety of analysis solutions and tools and hosts a large amount of relevant data, with more being submitted and released on a regular basis. Galaxy on AnVIL and its Toolshed allow for users to utilize even more tools with specific versions. Due to the lower knowledge/comfort score associated with Galaxy on AnVIL, this could be a potential area to prioritize in developing training materials so that users are empowered to bring their tools to the AnVIL if they are not already/directly supported.

The vast majority of users in our sample preferred virtual training opportunities over the other suggestions. The preference for virtual training could be due to the geographic and institutional spread of the AnVIL Community, funding constraints for travel, or any of the benefits that virtual events offer such as flexibility or increased accessibility for those with disabilities [49,50]. This poll was conducted before the inaugural AnVIL Community Conference and future polls and conversation may show this to be a preferred option as well. The AnVIL Outreach Team supports virtual training opportunities through several modalities (workshops, demonstrations, a 24/7 community support forum, webbooks, YouTube shorts, etc.). Users were largely aware of these support opportunities (namely AnVIL Demos and the support forum), even if utilization was lower. Training opportunities like workshops typically cover associated cloud computing costs to reduce the cost barrier. Handling this barrier for truly asynchronous training opportunities may be another area to prioritize. A solution could include providing estimates for the cloud computing cost of completing the training and an appendix instructing learners how to review incurred costs.

The third aim of this poll was to assess researchers’ technological comfort with cloud-based genomic analysis tools. Users appeared to be fairly comfortable with the cloud and running interactive analyses on the cloud, suggesting barriers to adoption could be platform specific rather than reluctance to use cloud tools. Less comfort or knowledge was observed with topics such as containers, workflows, and interactive analysis with Galaxy on AnVIL. In addition the TDR and DUOS systems had a low level of reported comfort. These are all areas where documentation or training could be advertised or further developed.

The final aim of this poll was to identify computational and data analysis resource needs. Several of the findings related to this aim have been discussed already – perhaps the most consequential being AnVIL’s support for analysis tools and acting as a data repository where researchers can store and access data. An additional conclusion relates to the future computational needs of AnVIL users. Most returning users reported that they foresee needing large amounts of storage in the future. This is unsurprising due to the vast amounts of data generated in the biomedical research field [39]. The responses and observations within this aim suggest that AnVIL is providing the computational and data analysis resource needs of users whether it be enabling data transfer from high performance computers, providing a variety of data analysis tools, or hosting popular datasets and enabling access as appropriate.

The findings of this poll are a snapshot from a limited set of current and potential AnVIL users. Responses may be biased by a variety of known phenomena, such as central tendency bias [51] when rating their knowledge or comfort with various technologies, or a non-response bias where respondents feel more strongly than those who did not respond [52–55]. In constructing the poll, care was taken to avoid common pitfalls in question design and poll administration that have been found to introduce or intensify bias [56,57]. We do not make statistical comparisons between groups due to the small sample size. While the number of responses we received was lower than expected or preferred, we view this poll and analysis as a worthwhile exercise in data gathering and a starting point where we make broad observations that the AnVIL Team then brings into conversations going forward. The data collection and analysis have helped the AnVIL Team to learn more about the AnVIL user base, previewing their needs and preferences. This in turn benefits researchers and clinicians who are interested in coming to the AnVIL so that they can see what the community currently looks like, perhaps even enabling them to find like-minded collaborators.

While we did not observe returning AnVIL users working within industry-based institutions in our sample, by considering other sources of user information (e.g., publications referencing the use of AnVIL or username domains), we observe that some users of the AnVIL work with industry-based institutions/companies. It is possible our sample did not capture that user base because of how the poll was disseminated. More work may be needed to understand the work and needs of AnVIL’s industry-based partners.

We would like to repeat this poll in the future, tracking how responses change over time and using the responses as a starting point for community discussion and more in-depth user interviews [58]. For future iterations of the poll, some adaptations may be helpful in order to target different user bases (such as data submitters or administrators that are supporting projects but not necessarily running analyses) with specialized questions/poll subparts rather than using a single set of questions. Future iterations or user interviews should additionally explore why users are motivated to use the AnVIL. With regards to user interviews, because the AnVIL Community is relatively new and there is little established knowledge about the user base, it is possible that in-depth qualitative data from user interviews will lead to more novel and intricate insights [59]. Though conclusions from user interviews will be even less generalizable than this small-sample poll, and researcher bias may impact how the conversation is steered or what parts of the conversation are highlighted later.

We are committed to transparency in sharing the poll, its analysis, and our findings. Users may be more comfortable or prioritize making requests or voicing a need/opinion if they see others have the same thoughts [60,61]. The AnVIL portal points to places where users can get help, ask questions, or provide feedback: https://anvilproject.org/help. Terra additionally displays active feature requests that can be explored: https://support.terra.bio/hc/en-us/community/topics/360000500452-Active-Feature-Requests. The findings of the poll will be useful in soliciting additional/future feedback from the AnVIL Community and will inform future platform development as well as enhance user support strategies. The AnVIL Team hopes to ultimately accelerate the adoption of cloud-based genomic analysis technologies and grow the AnVIL Community.

## Acknowledgements

This work was carried out under National Institutes of Health, National Human Genome Research Institute Project Number 5U24HG010263.

