## Supplemental Table 1 for "Insights into the Datasets, Tools, and Training Needs of the AnVIL Community: 2024"

| | | Examine the background and current work of users to develop appropriate personas | Understand barriers to platform adoption and user preferences for training and support | Assess researchers' technological comfort with cloud-based genomic analysis tools | Identify computational and data analysis resource needs |
|---|---|---|---|---|---|
| Part 1: Self-identify | | *x* | | | |
| | How would you describe your current usage of the AnVIL platform? | *x* | | | |
| Part 2: Feature importance + Returning user specific Q's | | *x* | *x* | *x* | *x* |
| | Rank features according to importance | | *x* | | *x* |
| | Knowledge of tech/data resources on the AnVIL | | | *x* | |
| | Length of use | *x* | | | |
| | Favorite AnVIL feature | *x* | | | |
| | Foreseeable computational/storage needs | | | | *x* |
| | Recommendation likelihood | *x* | | | |
| Part 3: Demographics | | *x* | | | |
| | Highest Degree | *x* | | | |
| | Industry | *x* | | | |
| | Kind of work | *x* | | | |
| | Institutional affiliation | *x* | | | |
| | Consortia affiliations | *x* | | | |
| Part 4: Experience | | *x* | | *x* | *x* |
| | Tech/data resource knowledge separate from the AnVIL | | | *x* | |
| | Types of data analyzed | | | | *x* |
| | Experience with human clinical, human genomic, or non-human genomic data | *x* | | | |
| | General & specific interest in controlled access datasets | | | | *x* |
| Part 5: Awareness | | | *x* | | |
| | Monthly AnVIL Demos | | *x* | | |
| | AnVIL Support | | *x* | | |
| Part 6: Preferences | | | *x* | *x* | *x* |
| | Rank various training workshop modalities | | *x* | | |
| | Where analyses are currently run | | | *x* | *x* |
| | DMS compliance/data repositories | | | | *x* |
| | Source of funds for cloud computing | | *x* | | |

Supplemental Table 1

**Relation of study aims to the design of the State of the AnVIL 2024 Community Poll (broken down by section and question).**

*Sections of the State of the AnVIL 2024 Community Poll (Part 1, Part 2, etc.) are listed in the first column as the row names. Column 2 provides the questions for each part. The rest of the column names are the enumerated study aims. X's are added at the intersection of any questions relevant to a particular study aim.*
