## Supplemental Table 2 for "Insights into the Datasets, Tools, and Training Needs of the AnVIL Community: 2024"

| Responses | Support Option | Awareness | Use (Past or Future) | Use (Past only) Demo: "Attend" |
|---|---|---|---|---|
| No, didn't know of | Demo | | | |
| | Support Forum | | | |
| No, but aware of | Demo | x | | |
| | Support Forum | x | | |
| Not yet, but am registered to | Demo | x | x | |
| Yes, one | Demo | x | x | x |
| Yes, multiple | Demo | x | x | x |
| Answered someone's post | Support Forum | x | x | x |
| Posted in | Support Forum | x | x | x |
| Read through other's posts | Support Forum | x | x | x |

Supplemental Table 2

**Recategorization of AnVIL Demo and Support Forum Utilization Raw Responses into Awareness and Use**

*This table provides all possible responses to the two questions asking respondents about their utilization of the AnVIL Demos and Support Forum. The second column describes which responses are applicable for each support option. The last 3 columns use an x if the raw response aligns with awareness or use.*
